# Metabolomic investigation of the pseudouridimycin producer, a prolific streptomycete

**DOI:** 10.1101/2020.11.05.369249

**Authors:** Marianna Iorio, Sahar Davatgarbenam, Stefania Serina, Paolo Criscenzo, Mitja M. Zdouc, Matteo Simone, Sonia I. Maffioli, Richard H. Ebright, Stefano Donadio, Margherita Sosio

## Abstract

We report a metabolomic analysis of *Streptomyces* sp. ID38640, a soil isolate that produces the bacterial RNA polymerase inhibitor pseudouridimycin. The analysis was performed on the wild type and on ten different *pum* mutants blocked at different steps in pseudouridimycin biosynthesis. The results indicate that *Streptomyces* sp. ID38640 is able to produce, in addition to pseudouridimcyin, lydicamycins and deferroxiamines, as previously reported, also the lassopeptide ulleungdin, the non-ribosomal peptide antipain and the osmoprotectant ectoine. The corresponding biosynthetic gene clusters were readily identified in the strain genome. We also detected the known compound pyridindolol, for which we propose a previously unreported biosynthetic gene cluster, as well as three families of unknown metabolites. Remarkably, the levels of the different metabolites varied strongly in the different mutant strains, allowing detection of metabolites not normally seen in the wild type. Three newly constructed *pum* mutants, along with systematic investigation of the accumulated metabolites, shed further lights on pseudouridimycin biosynthesis. We also show that several *Streptomyces* strains, harboring the *pum* biosynthetic gene cluster and unrelated to ID38640, readily produce pseudouridimycin.

## INTRODUCTION

After several decades of intensive screening, natural products still represent the best source of life-saving drugs, such as antibacterial and antitumor compounds.^1^ These specialized metabolites, as natural products are often referred to, have multiple biological activities that have become useful for human well-being. However, they can also play important roles in cell-cell communication or nutrient uptake.^2^ Specialized metabolites produced by soil-dwelling bacteria are no exception.

Several microbial genera belonging to different orders within the phylum Actinobacteria, commonly called actinomycetes, hold the genetic information for the synthesis of numerous specialized metabolites, devoting up to 10% of their genomes to this trait. The availability of multiple genome sequences and a variety of analysis tools such as antiSMASH, PRISM and BIG-SCAPE/CORASON allow the rapid identification of biosynthetic gene clusters (BGCs) in bacterial genomes.^3–5^ While genomic analyses are progressing fast, the majority of BGCs remain experimentally uncharacterized and yet to be associated to the cognate specialized metabolites. Different methods have been explored to harness such large biosynthetic potential, including cultivating strains in the presence of elicitors or stress substances,^6,7^ promoter refactoring,^8^ manipulating pathway specific regulators^9^ or using transcription factor decoys.^10^ Whatever the approach, detection of metabolites heavily relies on liquid chromatography (LC) coupled to mass spectrometry (MS). Significant advances have been made in metabolomics with new tools for organizing and analyzing MS data along with useful databases, making “omics” approaches very useful for prioritizing strains or molecules for further investigations.^11–13^

*Streptomyces* sp. ID38640, isolated from an Italian soil sample, is the producer of pseudouridimycin (PUM), the first nucleoside analog inhibitor of bacterial RNA polymerase, endowed with promising activity against Gram-positive and Gram-negative bacteria.^14^ PUM belongs to a small family of *C*-nucleosides antibiotics, which includes formycin, malayamycin and ezomycin.^15, 16^ We have previously analyzed the PUM pathways through knockouts of several *pum* genes present within the PUM BGC, leading to the first biosynthetic pathway for a *C*-nucleoside antibiotic.^17^ This work has also shown that the pseudouridine synthase PumJ, a key biosynthetic enzyme in the PUM pathway, is present in diverse, taxonomically unrelated microorganisms, suggesting a widespread distribution of yet-to-be-discovered *C*-nucleoside antibiotics.^17^

Blocking PUM biosynthesis in the producer strain *Streptomyces* sp. ID38640 led to altered production of the siderophore desferroxiamine and of the polyketide lydicamycin, two unrelated specialized metabolites.^17^ Here, we extend these findings by a systematic evaluation of MS profiles in available and *ad hoc* generated *pum* mutants, correlating metabolites with the corresponding BGCs. Overall, we were able to establish seven metabolite-BGC pairs, including proposing a new BGC for a known metabolite. We also show that PUM can be easily detected in *Streptomyces* strains harboring the *pum* BGC. Finally, our analyses provide additional insights into the PUM biosynthetic pathway.

## RESULTS and DISCUSSION

### Metabolomic analysis of pum mutants

Following earlier observations,^17^ we deemed interesting to globally analyze the metabolome of *pum* knockout mutants. To this end, the wild type (WT) strain, along with the knockout mutant strains listed in Table 1, all unable to produce PUM but accumulating different intermediates, were cultivated in two different media and analyzed by LC-MS at 24-hour intervals over four days. Samples were analyzed after solvent extraction of the whole culture (FE samples) and by direct analysis of the cleared broth (SN samples), which allowed detection of hydrophilic metabolites. The LC-MS/MS data from 44 samples, derived from 11 strains cultivated in two different media with two samples per culture, were subjected to the Global Natural Products Social (GNPS) Molecular Networking analysis^11^ and visualized using Cytoscape.^18^ This procedure clusters together spectra with identical MS^2^ pattern, forming the nodes represented by rectangles in Figure 1. Each fragmentation pattern is then pairwise compared with all the others and the ones sharing at least 4 fragments are connected with a line (dubbed cosine) forming a network. Each network represents a family of potentially related metabolites. Analysis with Cytoscape provides an intuitive visualization of metabolite distribution according to strain, medium or sample.

**Table 1.**
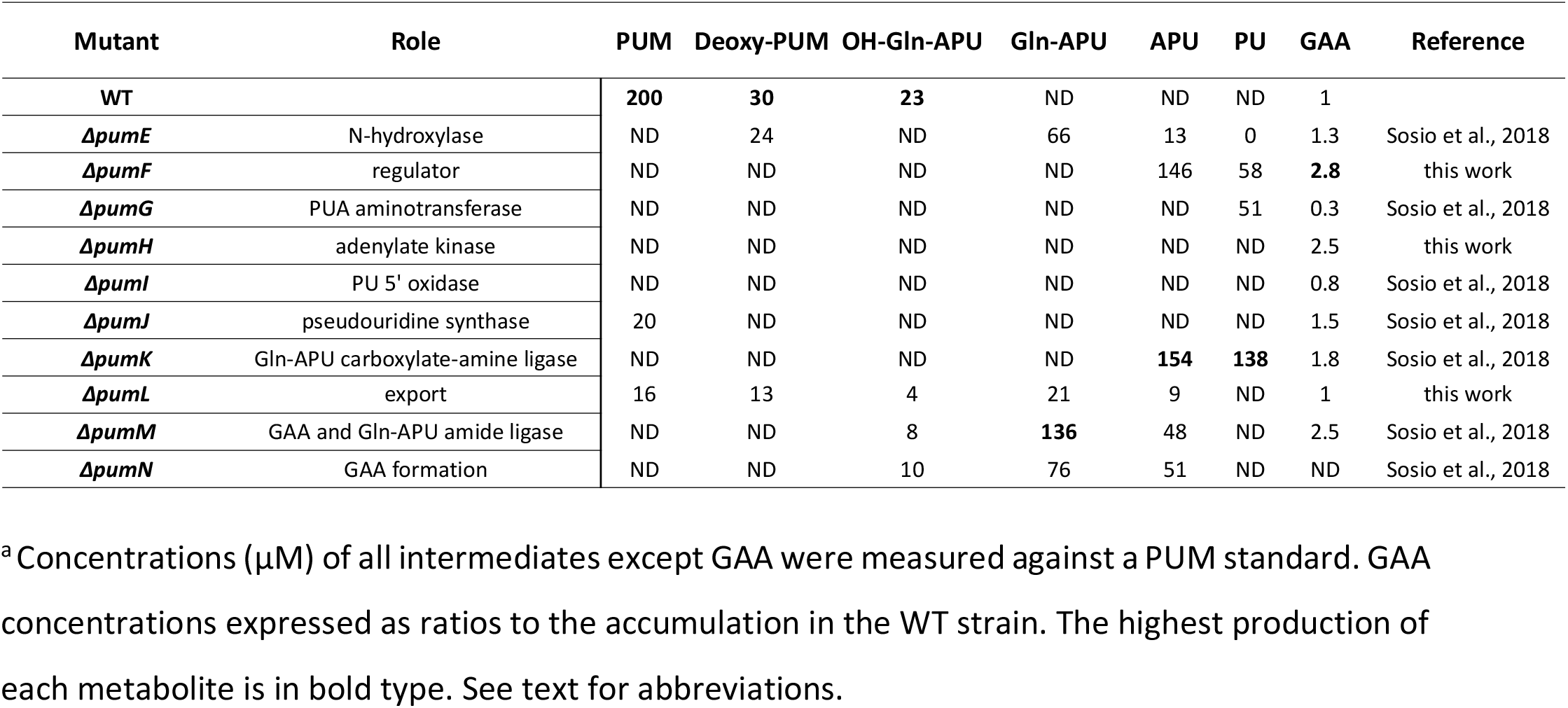
*pum* mutants and accumulated intermediates.^a^

| Mutant | Role | PUM | Deoxy-PUM | OH-Gln-APU | Gln-APU | APU | PU | GAA | Reference |
| --- | --- | --- | --- | --- | --- | --- | --- | --- | --- |
| **WT** |  | **200** | **30** | **23** | ND | ND | ND | 1 |  |
| *ΔpumE* | N-hydroxylase | ND | 24 | ND | 66 | 13 | 0 | 1.3 | Sosio et al., 2018 |
| *ΔpumF* | regulator | ND | ND | ND | ND | 146 | 58 | **2.8** | this work |
| *ΔpumG* | PUA aminotransferase | ND | ND | ND | ND | ND | 51 | 0.3 | Sosio et al., 2018 |
| *ΔpumH* | adenylate kinase | ND | ND | ND | ND | ND | ND | 2.5 | this work |
| *ΔpumI* | PU 5' oxidase | ND | ND | ND | ND | ND | ND | 0.8 | Sosio et al., 2018 |
| *ΔpumJ* | pseudouridine synthase | 20 | ND | ND | ND | ND | ND | 1.5 | Sosio et al., 2018 |
| *ΔpumK* | Gln-APU carboxylate-amine ligase | ND | ND | ND | ND | **154** | **138** | 1.8 | Sosio et al., 2018 |
| *ΔpumL* | export | 16 | 13 | 4 | 21 | 9 | ND | 1 | this work |
| *ΔpumM* | GAA and Gln-APU amide ligase | ND | ND | 8 | **136** | 48 | ND | 2.5 | Sosio et al., 2018 |
| *ΔpumN* | GAA formation | ND | ND | 10 | 76 | 51 | ND | ND | Sosio et al., 2018 |

<sup>a</sup> Concentrations (μM) of all intermediates except GAA were measured against a PUM standard. GAA concentrations expressed as ratios to the accumulation in the WT strain. The highest production of each metabolite is in bold type. See text for abbreviations.

**Figure 1.**
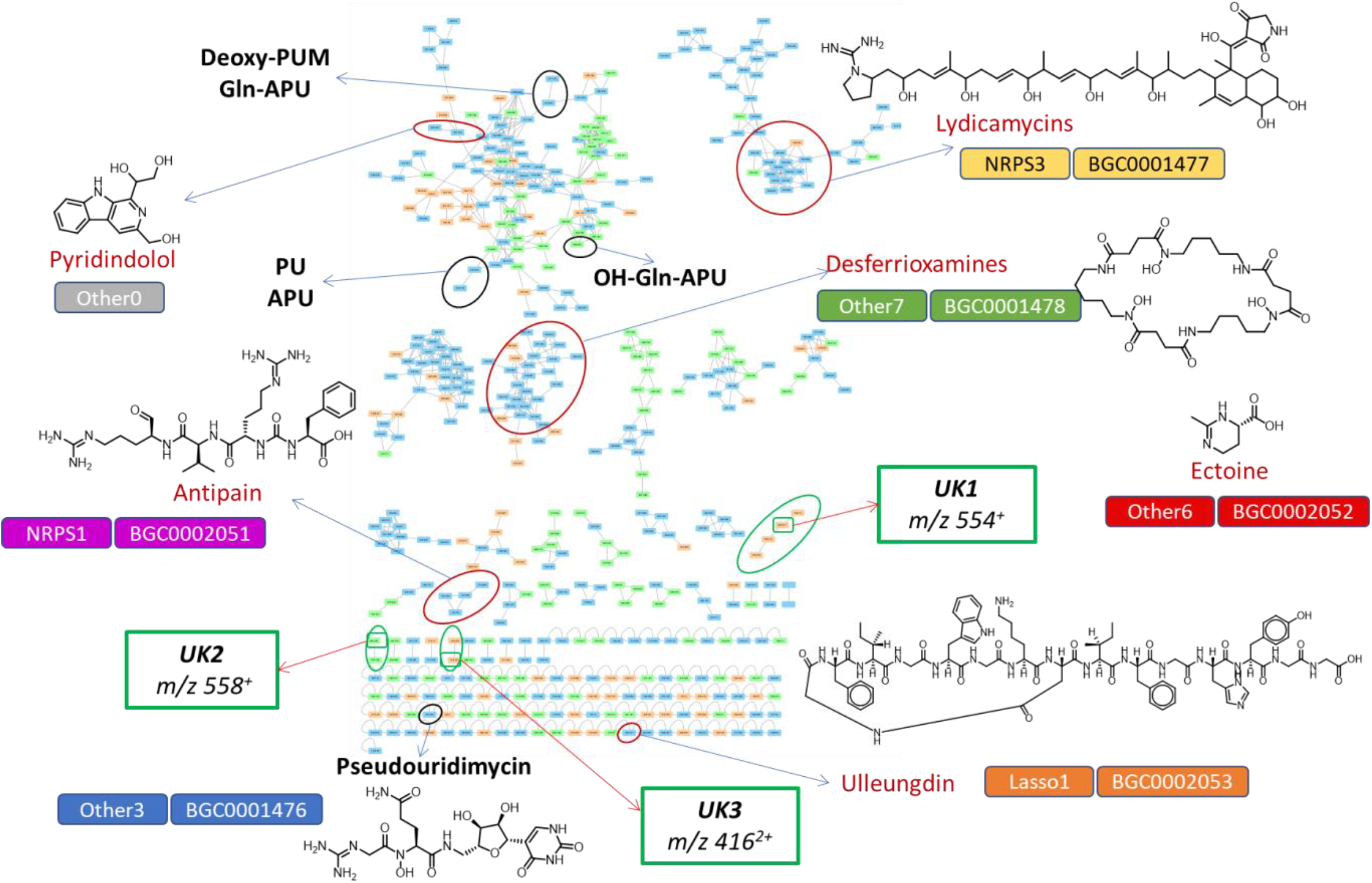
Molecular network of samples from *Streptomyces* sp. ID38640 and ten knockout *pum* mutants. The analysis includes two cultures from each strain and two samples per culture. Node colors give the contributing medium: orange for M8, green for PumP1, light blue for nodes observed in both media. Black circles indicate PUM-related nodes, red circles indicate nodes corresponding to known compounds, green circles show unknown metabolites. The associated BGCs, as listed in Table 2, are shown next to each metabolite.

The molecular network of Fig. 1 (derived from 72-h samples) consists of 475 features, including media components, 369 of which are organized in 36 molecular families, while the remaining 106 features are singletons. Similar results were observed when analyzing samples at different time points, but 72-h samples appeared as those with the highest metabolite richness. The two tested media were afforded equivalent numbers of signals, each showing about 20% unique signals. All strains contributed with 200–300 signals each, with *ΔpumH* and *ΔpumK* being at the bottom and top end, respectively. Similarly, 50% of signals were shared between the FE and SN samples, with 25% unique signals contributed by each sample. This might be surprising since the FE samples is expected to represent the entire metabolome. However, the solvent present in the FE samples pushes hydrophilic molecules to the solvent front during chromatography, thus preventing detection of these metabolites.

**Table 2.**
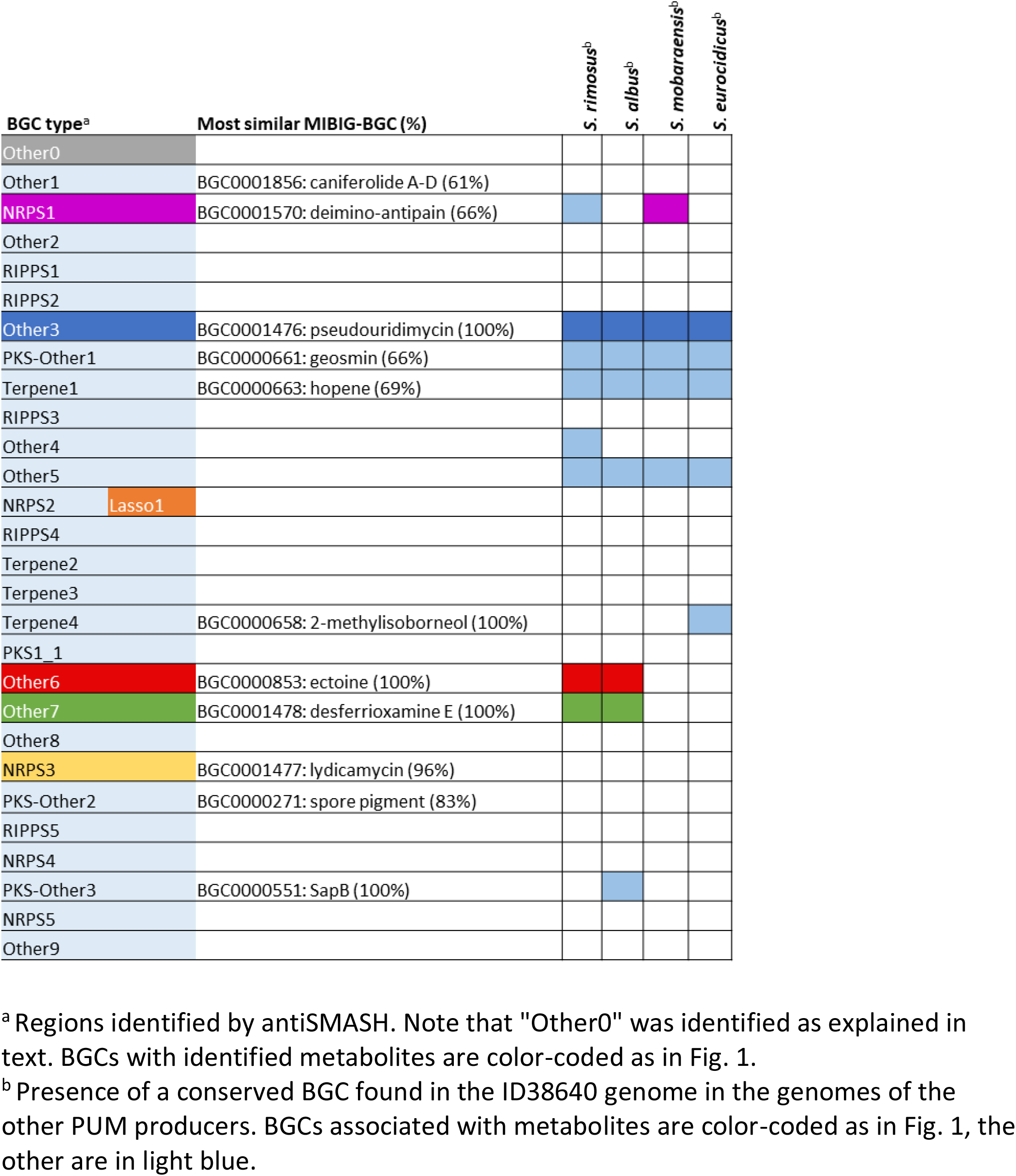
BGCs identified in ID38640.

### Correlating metabolites with biosynthetic gene clusters

From a draft genome of *Streptomyces* sp. ID38640, antiSMASH revealed 27 distinct regions harboring BGCs and a predicted metabolic diversity including five non-ribosomal peptides, four polyketides, six ribosomally synthesized postranslationally modified peptides, four terpenes, as well as at least nine regions classified as “other” (Table 2). Overall, 11 BGCs find a match with related BGCs in the MIBiG database.

Consistent with previous work,^17^ we have detected molecular networks corresponding to the lydicamycins and desferrioxamines in all tested samples from both fermentation media (Figure 1, Table 3). In particular, highest amounts of lydicamycins and desferrioxamines were detected in the *ΔpumI* and *ΔpumN* mutants, respectively (Table 3; Supplementary Fig. 1).

**Table 3.** Relative amounts of the identified metabolites in the different *pum* mutants. Amounts are expressed as ratios to those observed in the WT strain in the same medium. Note that for UK1 through UK3, which are not detected in the WT, their presence is indicated with an “X”. The highest relative amounts of metabolites are in bold type.

| Mutant | Major accumulated PUM metabolite(s) | Desferoxamines | Pyridindolol | Ectoine | Antipain | Ulleungdin | Lydicamycins | UK1 | UK2 | UK3 |
| --- | --- | --- | --- | --- | --- | --- | --- | --- | --- | --- |
| WT | PUM | 1.0 | 1.0 | 1.0 (traces) | **1.0** | 1.0 (traces) | 1.0 | ND | ND | ND |
| *ΔpumE* | Gln-APU | 2.0 | 0.8 | 9.0 | ND | **20.0** | 0.1 | X | ND | ND |
| *ΔpumF* | APU | 1.8 | 0.9 | 5.4 | 0.6 | 1.0 | 1.8 | ND | ND | ND |
| *ΔpumG* | PU | 1.3 | ND | 5.4 | 0.8 | 1.0 | 0.1 | **X** | **X** | ND |
| *ΔpumH* | GAA | 2.9 | ND | 7.4 | ND | 7.0 | 0.1 | ND | ND | ND |
| *ΔpumI* | GAA | 1.7 | 1.2 | 1.0 | ND | 3.0 | **3.2** | ND | ND | ND |
| *ΔpumJ* | GAA | 2.5 | ND | 9.6 | 0.9 | 1.0 | 0.4 | ND | ND | ND |
| *ΔpumK* | PU and APU | 2.9 | 1.4 | **10.6** | 0.1 | 5.0 | 1.2 | ND | ND | ND |
| *ΔpumL* | Gln-APU | 2.1 | ND | 2.0 | 0.6 | 1.0 | 0.1 | ND | X | **X** |
| *ΔpumM* | Gln-APU | 2.7 | 0.6 | 4.4 | 0.9 | 1.0 | 0.5 | X | ND | ND |
| *ΔpumN* | Gln-APU | **3.3** | **2.5** | 3.0 | 0.4 | 2.0 | 1.2 | X | ND | ND |
| Best medium |  | M8 | PumP1 | PumP1 | Both | M8 | Both | M8 | PumP1 | M8 |

The molecular network includes a family with a node at *m/z* 303 [M+2H]^2+^ (Fig. 1) whose HR-MS fragmentation pattern and UV spectrum (Supplementary Fig. 2) matched those of an authentic standard within this family, corresponding to the ureylene-containing oligopeptide antipain,^19^ produced by non-ribosomal peptide synthetases in numerous bacteria and acting as a protease inhibitor.^20^ Accordingly, we found the corresponding BGC in the ID38640 genome, showing 61-78% identical genes with the related BGCs in the MIBIG database^21^ (Table 2). The antipain molecular family has been found in the WT strain and in most *pum* mutants in both media, with highest amounts detected in the WT (Table 3; Supplementary Fig. 1).

The *Streptomyces* sp. ID38640 genome contains a lassopeptide BGC (Table 2; Supplementary Fig. 3) and the predicted core peptide is identical to ulleungdin, a recently reported lassopeptide from *Streptomyces* sp. KCB13F003.^22^ Consistently, we found a self-loop feature (Fig. 1) corresponding to an exact mass of 796.8835 [M+2H]^2+^ (ulleungdin calculated mass 796.8859 [M+2H]^2+^). The identity of this metabolite with ulleungdin was confirmed by the MS fragmentation pattern (Supplementary Fig. 3). The lassopeptide has been detected in all tested strains irrespective of the fermentation medium, with several *pum* mutants producing sizeable levels in contrast with the trace amounts observed in the WT (Table 3; Supplementary Fig. 1).

The molecular networking analysis of Fig. 1 was carried out using a cosine score above 0.7. This value was also used to form MS clusters. This filter excluded ectoine, a methyl, tetrahydropyrimidinecarboxylic acid that protects many bacterial species from osmotic stress, since this metabolite shows very poor fragmentation. Nonetheless, we were able to identify a peak at 1.1 min, consistent with ectoine hydrophilicity, with a matching exact mass (found *m/z* 143.0813 [M+H]^+^, calculated *m/z* 143.0815 [M+H]^+^; Supplementary Fig. 4). Ectoine has been detected in all tested strains (Table 3; Supplementary Fig. 1) and the corresponding BGC, a common encounter in *Streptomyces* genomes, is present in the ID38640 genome (Table 2).

The molecular network of Fig. 1 highlighted three additional families corresponding to unknown molecules. Four different *pum* mutants, when grown in M8 medium only, produced four related species with *m/z* 536 [M+H]^+^, *m/z* 554 [M+H]^+^, *m/z* 555 [M+H]^+^ and *m/z* 642 [M+H]^+^, named “UK1” (Table 3; Supplementary Fig. 5). Two additional, related signals (*m/z* 558 [M+H]^+^ and *m/z* 530 [M+H]^+;^ “UK2” in Fig. 1) were observed in samples from the *ΔpumG* and *ΔpumL* strains, only when grown in PumP1 medium (Table 3; Supplementary Fig. 5). Moreover, two signals corresponding to double charged masses *m/z* 416 [M+2H]^2+^ and *m/z* 423 [M+2H]^2+^ were observed only in samples from *ΔpumL* in M8 medium (“UK3”, Fig. 1; Table 3; Supplementary Fig. 5). No matches of these signals were found in the Dictionary of Natural Products and in the Natural Product Atlas,^27, 28^ while our internal set of 5,200 *Streptomyces* molecular fingerprints indicated that UK1, UK2 and UK3 are rare occurrences. Thus, these molecules may represent novel metabolites worthy of further investigations.

We also observed a molecular family consisting of *m/z* 259 [M+H]^+^ and of *m/z* 421 [M+H]^+^, with exact masses of *m/z* 259.1089 [M+H]^+^ and 421.1615 [M+H]^+^, respectively (Supplementary Fig. 6). The associated LC peaks showed a UV-Vis spectrum with maxima at 254, 304 and 370 nm. These properties matched those of pyridindolol and its glucoside, produced by *Streptomyces alboverticillatus* and *Streptomyces parvulus*, respectively.^23, 24^ The MS2 fragmentations are consistent with this annotation (Supplementary Fig. 6). These metabolites have been detected in almost all strains, with the *ΔpumN* mutant as the highest producer (Table 3; Supplementary Fig. 1).

Pyridindolol biosynthesis has not been studied so far and no BGC possibly linked to this metabolite could be found in the antiSMASH output. However, it has been reported that the β-carboline moiety present in pyridindolol is formed by a “Pictet-Spenglerase” (PSase), an enzyme joining an amino group with an aldehyde.^25^ While these enzymes are mostly encountered in plant alkaloid biosynthesis, the enzyme StnK2 has been shown to catalyze such a reaction during streptonigrin biosynthesis.^26^ Accordingly, we searched the ID38640 genome and identified QIK04791.1 as a likely PSase showing 49% sequence identity with StnK2 (Fig. 2; Table 4). The QIK04791.1-encoding region also specifies for a FAD-binding oxidoreductase, a long-chain fatty acid-CoA ligase, an aldehyde dehydrogenase, a histidine phosphatase and an F420-dependent oxidoreductase (Table 4). While most of these sequences have no paralogs in the streptonigrin BGC, a syntenic region with over 90% protein-to-protein identity (Fig. 2; Table 4) is present in the genome of the pyridindolol producer *Streptomyces alboverticillatus* (MUFU00000000). Accordingly, pyridindolol formation is predicted to require few steps: condensation of tryptophan with a C-3 unit, possibly glyceraldehyde(phosphate) by the PSase; aromatization of the newly formed ring by the FAD-binding- and/or the F420-dependent oxidoreductase; and reduction of the carboxyl group by the aldehyde dehydrogenase (Fig. 2). The order in which these hypothetical steps occur awaits experimental demonstration, as well as the possible participation of the conserved long-chain fatty acid-CoA ligase and histidine phosphatase present in the conserved segment. This BGC, which lies at one end of the genome sequence in *Streptomyces* sp. ID38640, has been added to Table2 and designated “Other0”.

**Table 4.** The proposed pyridindolol BGC.

| CDS | Size and Protein ID | Protein Family | Best MIBIG match, % identity (BGC) | Homolog<sup>a</sup> [strain, accession No., % identity] |
| --- | --- | --- | --- | --- |
| 1 | 326 aa<br>QIK04786.1 | Regulator |  | AraC family transcriptional regulator [*Streptomyces lydicus*, WP_127154781.1, 98%] |
| 2 | 406 aa<br>QIK04787.1 | flavin reductase |  | FAD-binding oxidoreductase [*Streptomyces lydicus*, WP_127154780.1 97%] |
| 3 | 574 aa<br>QIK04788.1 | flavin reductase |  | flavin reductase [*Streptomyces libani*, WP_159483585.1, 97%] |
| 4 | 94 aa<br>QIK04789.1 | hypothetical |  | lantibiotic dehydratase C-term region [*Streptomyces rimosus*, WP_125052965.1, 73%] |
| 5 | 90 aa<br>QIK04790.1 | hypothetical |  | Hypothetical prot [*Streptomyces lydicus*, AZS70204, 83%] |
| 6 | 322 aa<br>QIK04791.1 | Pictet-Spenglerase | StnK2, 49%<br>(BGC0001783) | hypothetical protein [*Streptomyces alboverticillatus*, WP_086571431.1, 95%] |
| 7 | 370 aa<br>QIK04792 | FAD-binding oxidoreductase | StnP2, 39%<br>(BGC0001783) | FAD-binding oxidoreductase [*Streptomyces alboverticillatus*, WP_086571429, 93%] |
| 8 | 352 aa<br>QIK10964.1 | long-chain fatty acid--CoA ligase |  | AMP-binding protein [*Streptomyces alboverticillatus*, WP_086571427.1, 95%] |
| 9 | 453 aa<br>QIK04793.1 | aldehyde dehydrogenase family protein |  | aldehyde dehydrogenase family protein [*Streptomyces alboverticillatus*, WP_086571425.1, 94%] |
| 10 | 218 aa<br>QIK10965.1 | histidine phosphatase family protein |  | histidine phosphatase family protein [*Streptomyces alboverticillatus*, WP_086571443.1, 87%] |
| 11 | 293 aa<br>QIK04794.1 | LLM class F420-dependent oxidoreductase |  | LLM class F420-dependent oxidoreductase [*Streptomyces alboverticillatus*, WP_086571441.1, 90%] |
| 12 |  | IS5/IS1182 family transposase |  |  |

Matches to the syntenic *S. alboverticillatus* region shown in Fig. 2 are in red.

**Figure 2.**
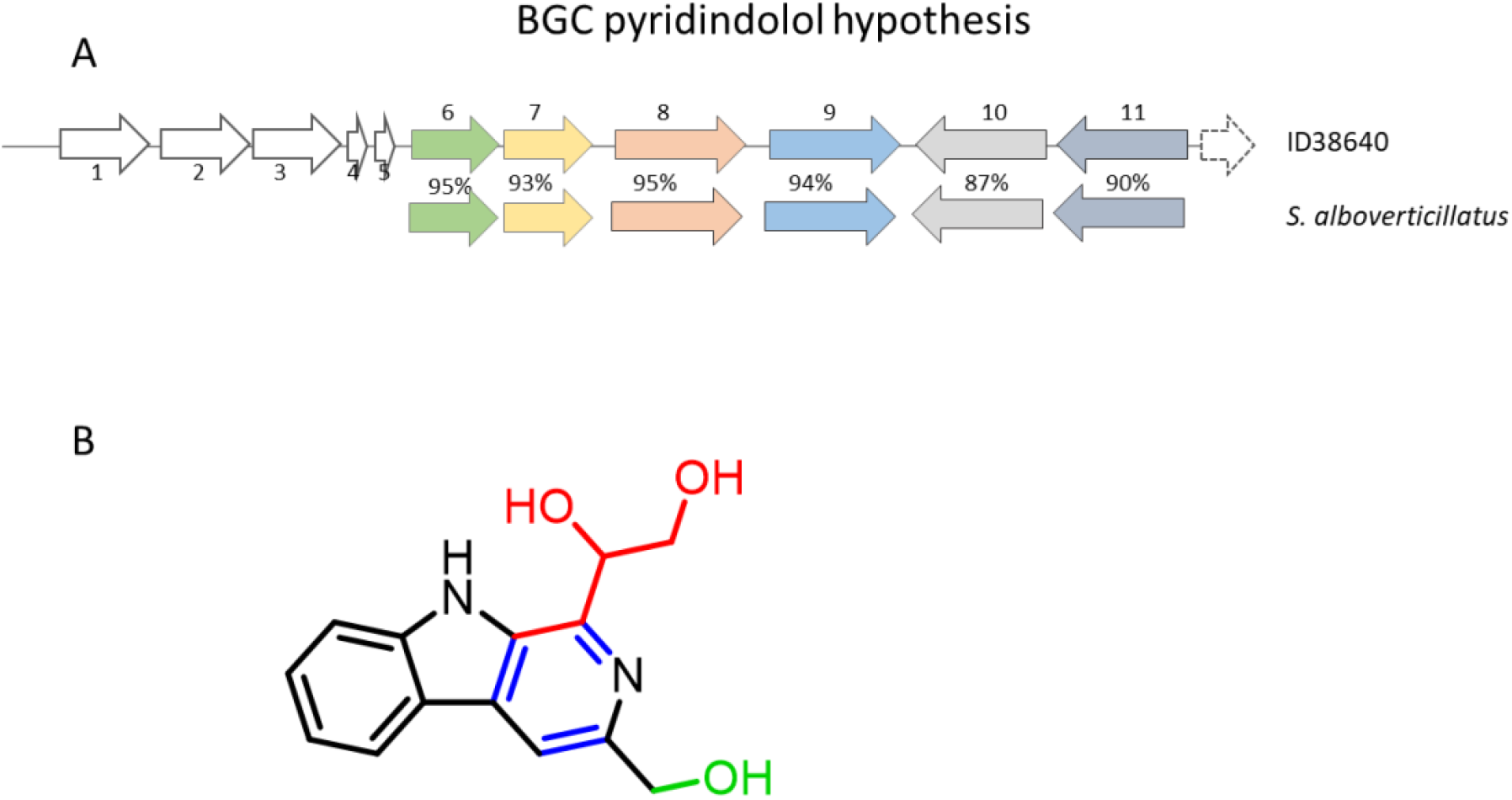
Putative pyridindolol BGC and hypothetical roles in metabolite formation. A) Identified region containing the PSase homolog from the ID38640 genome (top) and syntenic region found in the pyridindolol producer *S. alboverticillatus* (bottom). The percent identities between orthologs are shown, while deduced functions are reported in Table 4. B) Pyridindolol structure, highlighting the likely building blocks tryptophan and glyceraldehyde, and the modifications necessary for the final structure.

Overall, our work led to the identification of seven metabolite-BGC pairs (Table 2). This leaves 21 BGCs orphan of their product. This list includes BGCs for geosmin and methylisoborneol, volatile metabolites unlikely to be detected under our conditions, and hopene, unlikely to be present in our samples because of its lipophilicity. In total 18 BGCs await matching metabolites and 3 identified metabolites are still looking for a matching BGC.

Overall, this work has demonstrated that small changes in the genome facilitate the detection of additional metabolites. It has been previously reported that blocking biosynthesis of a specialized metabolite can facilitate detection of novel chemistry.^29^ However, we are not aware of studies showing that different blocks in a single pathway can significantly alter the metabolite levels of biosynthetically unrelated metabolites. Accumulation of a particular PUM intermediate does not appear the reason for altering the metabolic profiles: for example, ectoine levels are 10fold enhanced in the *ΔpumE, ΔpumJ* and *ΔpumK* mutants that accumulate different PUM intermediates (Table 1). At the same time, not all mutants accumulating the same PUM intermediate show similar increases in ectoine levels. The mechanism(s) leading to altered metabolic profiles are currently unknown and further work will be necessary to establish whether this phenomenon is an oddity of the PUM pathway. Nonetheless, the generation of distinct mutants from a single BGC might be useful not only for elucidating the corresponding biosynthetic pathway (see below), but also for altering regulatory circuits within the cell and allowing detection of chemistry hidden in wild-type strain.

### Additional insights into PUM biosynthesis

The functions of most of the *pum* genes and the molecular basis for PUM biosynthesis have been proposed from previous *in vivo* experiments and bioinformatic analyses,^17^ as summarized in Table 1. In this work, we generated knockout mutants in *pumF, pumH* and *pumL*. PumF shows 42-45% identity to SsaA and its orthologues NpsM and PacA, regulatory proteins present in the structurally related uridyl-peptide antibiotics sansanmycins, napsamycin and pacidamycin gene clusters, respectively.^30^ PumH, annotated as an adenylate kinase, shares 42% identity with PolQ2 and MalE from the polyoxin and malayamycin biosynthetic pathways, respectively.^31^ PumL shares 65% identity with a NocH-like protein belonging to the major facilitator superfamily.

Replacement of *pumF* with the apramycin resistance gene abolished PUM production and led to the accumulation of pseudouridine (PU), amino pseudouridine (APU) and guanidine acetate (GAA) (Table 1; Supplementary Fig. 7). These results indicate that the positive regulator PumF is necessary for antibiotic production and controls the conversion of APU into Gln-APU. Deletion of *pumL* resulted in very low yields of PUM and accumulation of several intermediates, consistent with its role in exporting the final pathway product (Table 1; Supplementary Fig. 7).

The phenotype of the *ΔpumH* mutant was more complex: it accumulated no PUM-related metabolite except for GAA (Table 1; Supplementary Fig. 7); and, unlike the *ΔpumJ* strain,^17^ PUM production could not be rescued by adding PU to the production medium. Thus, the *ΔpumH* phenotype was identical to that of the previously described *ΔpumI* mutant,^17^ which also accumulated no intermediate except for GAA and could not convert PU into PUM. The simplest interpretation of these results is that inactivation of *pumH* or *pumI* through insertion of the apramycin resistance gene has a polar effect on downstream genes. Consistently, *pumH* and *pumI*, and *pumI* and *pumJ*, have 20- and 4-bp overlaps, respectively, suggesting they belong to a single transcriptional unit. Insertion of the apramycin resistance cassette into the kinase-encoding *pumH* would thus disable also the oxidoreductase PumI and the pseudouridine synthase PumJ, while insertion into *pumI* would leave only *pumH* intact.

We also explored the PUM precursors accumulated in all the *pum* mutants in the molecular network. PUM itself (*m/z* 487 [M+H]^+^) appears as a single loop detected only in the WT and, to a lesser extent, in the *ΔpumJ* and *ΔpumL* mutants, irrespective of the cultivation medium (Figure 1 and Table 1). We also detected a molecular family including PU, *m/z* 244 [M+H]^+^, and APU, *m/z* 245 [M+H]^+^. PU accumulated only in the *ΔpumF, ΔpumG* and *ΔpumK* mutants, while APU was identified mainly in the *ΔpumF* and *ΔpumK* strains, with highest amount of both metabolites observed in *ΔpumK* (Table 1). The late PUM intermediates Gln-APU (*m/z* 372 [M+H]^+^) and deoxy-PUM (*m/z* 471 [M+H]^+^) cluster together (Figure 1). As expected, Gln-APU was found in samples from the *ΔpumE, ΔpumL, ΔpumN* and *ΔpumM* strains, the latter accumulating the highest amount. Deoxy-PUM was detected in samples from the WT, *ΔpumE* and *ΔpumL* strains, in agreement with previous results, with WT being the highest producer (Figure 1, Table 1). In a different portion of the same network we observed a signal at *m/z* 388 [M+H]^+^) consistent with *N*-hydroxy-Gln-APU (OH-Gln-APU), which corresponded to a hydrophilic peak with the pseudouridine-characteristic UV maximum at 263 nm. This species, which had been overlooked in previous work because of its low abundance (Table 1), was detected in samples from the WT and, in lower amounts, from the *ΔpumL, ΔpumM* and *ΔpumN* strains. The levels of the PU-containing intermediates sharing the same chromophore were quantified as reported in Table 1.

The results presented here allowed us to revise the previously proposed biosynthetic pathway for PUM (Fig. 3). During the early biosynthetic steps, the substrate for the kinase PumH might be uridine, PU or PU aldehyde (Fig. 3), with PumJ, PumI and PumG acting sequentially for *C-*isomerization, alcohol oxidation and amine formation, respectively. Eventually, the phosphate group is released by PumD or by housekeeping phosphatases. A key step in the pathway appears to be the conversion of APU into Gln-APU by PumK, as this conversion is controlled by the regulator PumF. The detection of OH-Gln-APU is consistent with the possibility that *N*-hydroxylation by PumE precedes GAA addition by PumM, consistent with the well-known facilitated hydroxylation of amines with respect to amides.^32^ In the absence of PumE, PumM can use Gln-APU as substrate, leading to deoxyPUM as a shunt metabolite. While in the absence of PumM, Gln-APU preferentially accumulates, suggesting that either conversion of Gln-APU into its hydroxyderivative is inefficient or expression of *pumE* is altered in such background. Finally, PumL appears to be a transporter for PUM.

**Figure 3.**
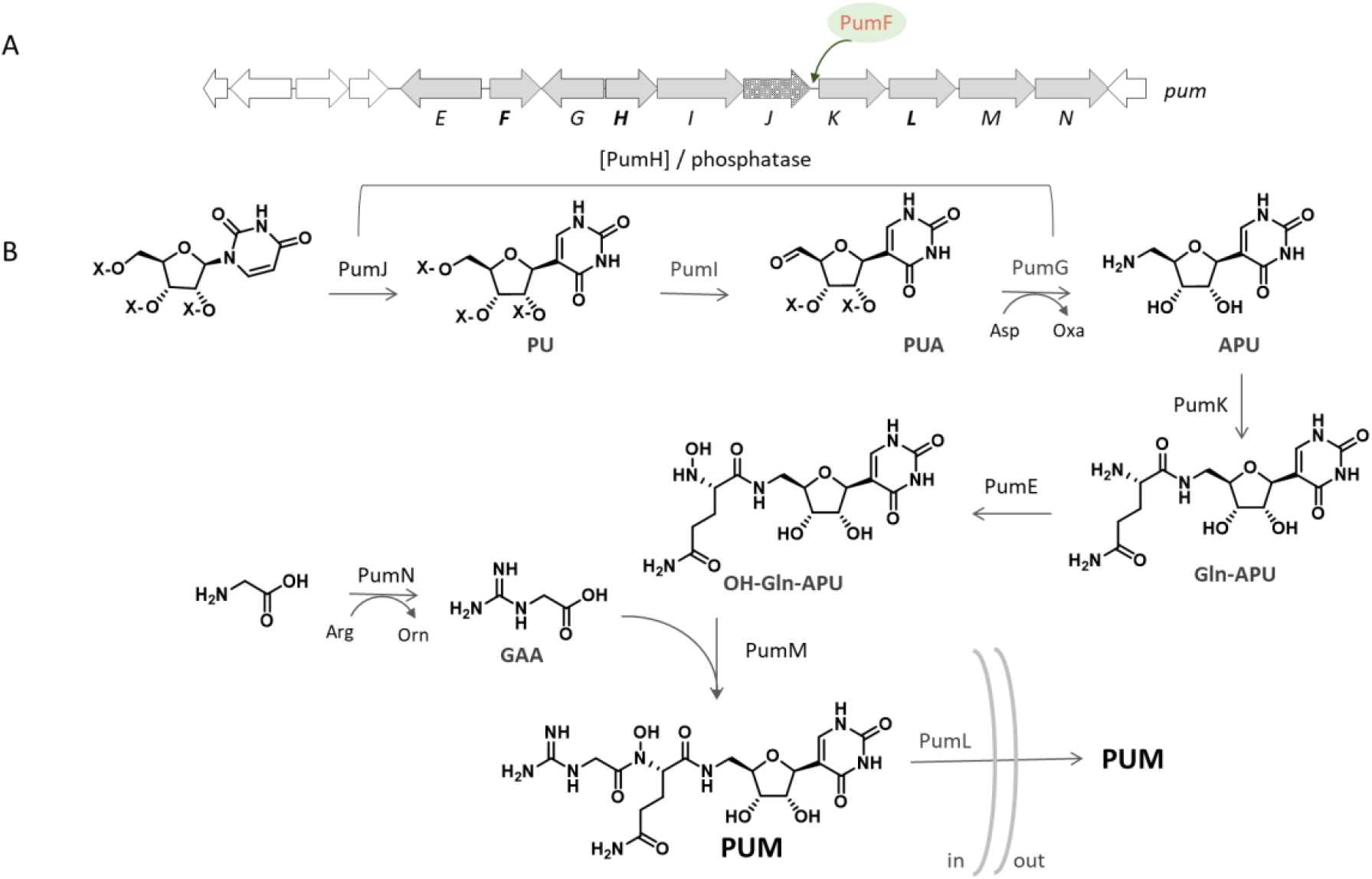
Revised biosynthetic pathway for pseudouridimycin. A) PUM BGC, with established role for PumF. B) proposed pathway. Enzymes and intermediates reported within brackets have not been experimentally determined. See text for abbreviations.

### A glimpse at other PUM producers

In previous work we have identified a number of related *pumJ*-like sequences in microbial genomes linked to putative BGCs,^17^ some of which were predicted to specify biosynthesis of PUM or closely related metabolites. In addition to a few strains from the NAICONS collection,^14^ PUM has been observed only in *Streptomyces albus* DSM 40763.^33, 34^ We thus chose a few *Streptomyces* strains that harbor a PUM BGC: *S. rimosus* ATCC 10970, the oxytetracycline producer; *S. mobaraensis* DSM 40847, the producer of the NADH reductase inhibitor piericidin; *S. eurocidicus* ATCC 27428, the producer of the antifungal polyene eurocidin; and *S. flocculus* DSM 40313, the producer of the aminoquinone antibiotic streptonigrin. [*S. flocculus* has been recently reclassified as later heterotypic synonyms of *S. albus* ^35^ and will be referred as *S. albus* DSM 40313 hereafter.] These compounds have been known for several decades and the producer strains have been investigated by several laboratories, but PUM production has to our knowledge not been observed.

These strains were grown in a single medium and analyzed at three different time points: LC-MS analysis of processed samples indicated that each *Streptomyces* species, in addition to the known metabolites oxytetracycline, piericidin, eurocidin D or streptonigrin, produced PUM, as indicated by a peak at 3.9 min with *m/z* 487 [M+H]^+^ corresponding to PUM (Figure 4a). In particular, *S. rimosus* and *S. mobaraensis* produced PUM at a level comparable to *Streptomyces* sp. ID38640 (around 200 μM), while *S. albus* and *S. eurocidicus* produced about 100 μM PUM. These results indicated that the PUM BGC in these species is actively expressed and that PUM can be easily detected when properly looked for.

**Figure 4.**
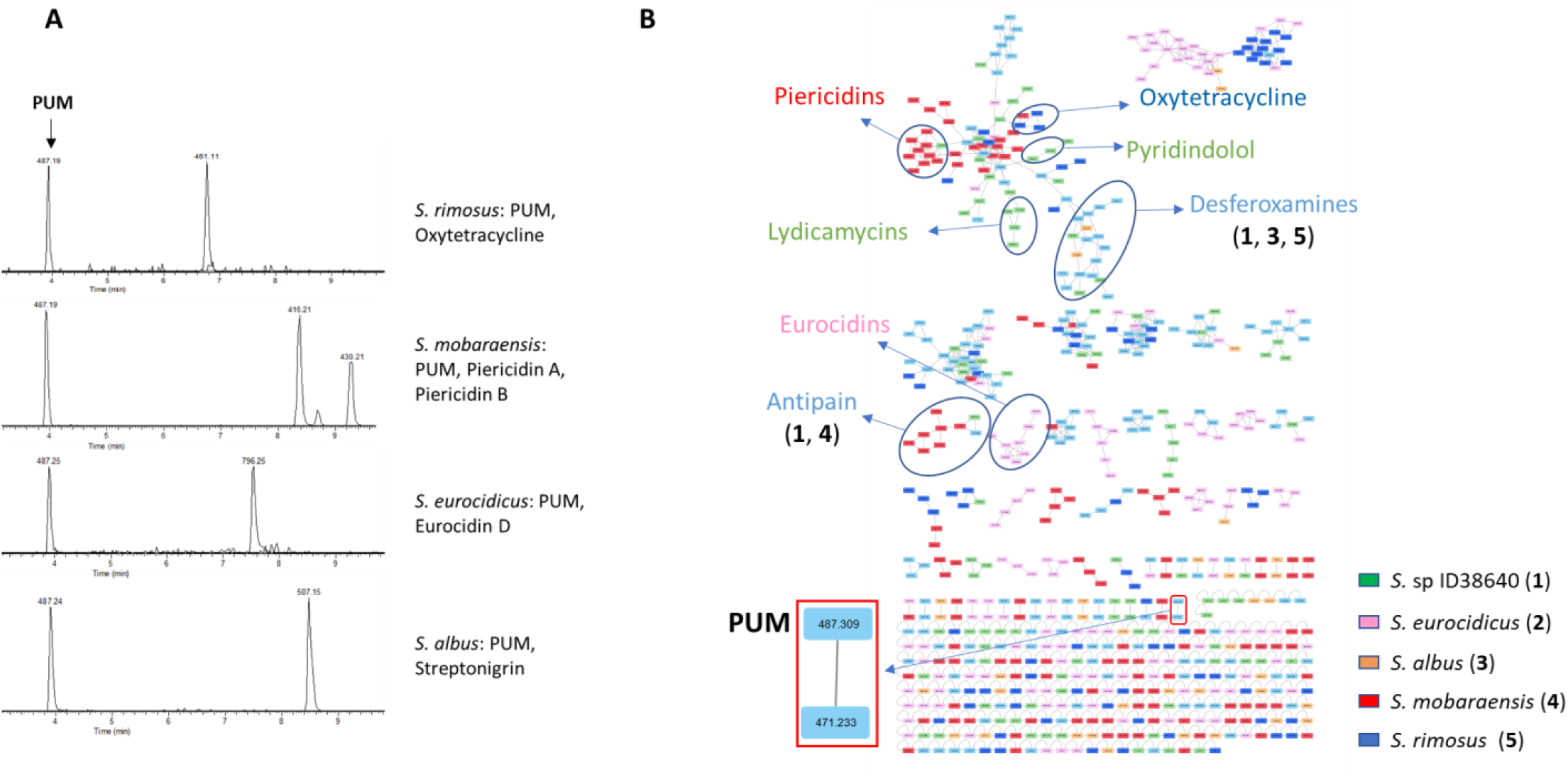
Analysis of PUM production in other *Streptomyces* strains. A) Extracted ion chromatograms of PUM (*m/z* 487 [M+H]+) and strain specific metabolites from *S. rimosus, S. mobaraensis, S. eurocidicus* and *S. albus*. B) Complete molecular network of two samples from each of the above PUM producers and from ID38640. Strain-specific features are color coded as: ID38640, green; *S. eurocidicus*, pink; *S. albus*, orange; *S. mobaraensis*, red; and *S. rimosus*, blue. Features detected in more than one strain are in light blue. All strains were cultivated in M8 medium and samples prepared at 72 hours.

To delineate the relationship of the different PUM producers, we resorted to autoMLST^36^ for a high-resolution species tree using multi-locus comparison. The resulting tree revealed ten major clades and three branches formed by a single strain each (Supplementary Fig. 8). *Streptomyces* sp. ID38640 belongs to clade 2 while, among the PUM producers reported above, only *S. rimosus* (clade 7) and *S. mobaraensis* (single-strain branch) were picked up. Next, we made a phylogenetic tree of the five PUM producers. Again, *Streptomyces* sp. ID38640 clusters together with *S. rimosus*, while *S. eurocidicus* and *S. mobaraensis* formed a separate clade (Supplementary Fig. 9). This analysis indicated that the PUM BGC is not restricted to a specific *Streptomyces* clade.

As reported in Table 2, the ID38640 genome does not harbor BGC for oxytetracycline, euricidin, piericidin or streptonigrin. We next investigated whether the other PUM-producing strains shared other metabolites with *Streptomyces* sp. ID38640, in a manner similar to the analysis of the different *pum* mutants described above. The resulting molecular network, represented in Figure 4b, contains 630 features, including media components, of which 385 (61%) are organized in 62 molecular families.

As highlighted by a red rectangle, a two-member family containing PUM and deoxy-PUM is found in samples from all strains. Of the families annotated in Fig. 1, lydicamycins, pyridindolol, ulleungdin and UK1 through UK3 remain ID38640-specific. Desferroxamines were seen in *S. rimosus* and *S. albus*, while antipain was detected in *S. mobaraensis*. Manual search, as explained above, showed that ectoine was present in extracts from *S. rimosus* and *S. albus*. As expected, the corresponding BGCs could be easily identified in the strains genomes (Table2). Several additional metabolites were dereplicated in the samples from the four other PUM producers, and the corresponding BGCs identified in the genomes (M.I., unpublished observations). However, none of these molecules matched any of the unannotated metabolites observed in *Streptomyces* sp. ID38640.

## CONCLUSIONS

*Streptomyces* sp. ID38640 has proved a prolific and versatile producer of different metabolites, many of which could become visible only after selective blocks in the PUM pathway. While we do not yet understand why production of unrelated metabolites is significantly enhanced in different *pum* mutants, the approach outlined here might be a simple way of catching two birds with a stone: on the one hand elucidating a biosynthetic pathway of interest, on the other hand looking for alterations in the metabolome. Possible targets for this approach might be the other PUM producers reported in this study. In any case, the work presented here, along with our previous studies,^37-38^ indicate that a metabolomic look at “old strains” can unveil previously overlooked chemistry, including novel metabolites. This sort of analyses will be undoubtedly facilitated by the growing Paired Omics Data Platform (https://pairedomicsdata.bioinformatics.nl).

### General Experimental Procedures

#### Bacterial strains and growth conditions

*Streptomyces* sp. ID38640, *S. flocculus* DSM 40313, *S. rimosus* ATCC 10970, *S. mobaraensis* DSM 40847, *S. eurocidicus* ATCC 27428 and the *pum* mutants were cultured as described.^14^ Briefly, mycelium from BTT plates was inoculated in 50-mL Erlenmeyer flask containing 15 mL of seed medium (20 g/L dextrose monohydrate, 2 g/L yeast extract, 8 g/L soybean meal, 1 g/L NaCl, and 4 g/L CaCO3, pH 7.3), and incubated 72 h at 28 °C. The production media were M8 ^39^ and PumP1 ^14^, which were inoculated with a 10% volume of the seed culture.

#### Construction of knockout mutants

The generation of *ΔpumF, ΔpumH* and *ΔpumL* strains followed described procedures,^17^ which involved amplification of two ∼1.0-kbp fragments (A and B) from genomic DNA using primers containing *Eco*RI and *Xba*I (fragment A) and *Xba*I and *Bam*HI (fragment B) tails (Supplementary Table 1), which were cloned into the *EcoRI-BamHI* sites of the vector pWHM3-*oriT-ΔXba*. In the resulting plasmid, the apramycin resistance gene was inserted at the *XbaI* site within the PCR-amplified *pum* segments to generate the knockout plasmid. The knockout plasmids were then introduced into *E. coli* ET12567/pUB307, whence they were conjugated into spores of *Streptomyces* sp. ID38640 as described.^17^ Double-crossover mutants were identified through PCR with diagnostic primers.

#### Genome sequence and bioinformatic analyses

Genome sequencing was performed by Cebitec Bielefeld University (Germany) using Illumina MiSeq / Genome Analyzer IIx / HiSeq 1000. BGCs were identified using the antiSMASH 5.0 at the default conditions.^3^ BLAST analysis of individual CDSs was performed against the MIBiG database of known BGCs ^21^ and against Protein Data Bank. Multilocus sequence analysis was performed with autoMLST in “denovo mode” and default settings.^36^

#### Samples for LC-MS analysis

For PUM-related metabolite analysis, 0.5 mL of the culture was centrifuged at 13200 rpm for 2 min and the supernatant was filtered through a 0.2-μm membrane (EuroClone), generating the SN sample. Full extracts (FE) were prepared by transferring a 0.5-mL sample from cultures into a 2-mL Eppendorf tube containing 0.5 mL MeOH. After 1 hour at 55 °C under constant shaking, the sample was centrifuged for 10 min at 13,200 rpm and the supernatant was recovered and transferred into a 1.5-mL glass vial.

#### Metabolite analysis

LC-MS analyses were performed with on a Dionex UltiMate 3000 coupled with an LCQ Fleet (Thermo scientific) mass spectrometer equipped with an electrospray interface (ESI) and a tridimensional ion trap. The column was an Atlantis T3 C18 5 mm x 4.6 mm x 50 mm maintained at 40 °C at a flow rate of 0.8 mL/min. Phases A and B were 0.05% trifluoroacetic acid in water and acetonitrile, respectively. SN samples were analyzed using the following gradient: 0 to 25% phase B in 4 min, followed by a 2-min wash at 90% and a 3-min re-equilibration at 0% phase B. The gradient used for FEs was a 14-min multistep program that consisted of 10, 10, 95, 95, 10 and 10% phase B at 0, 1, 7, 12, 12.5 and 14 min, respectively. UV-VIS signals (190-600 nm) were acquired using the diode array detector. The *m/z* ranges were set at 120-1500 and 200-2000 for SNs and FEs, respectively, with ESI conditions as follows: spray voltage of 3500 V, capillary temperature of 275 °C, sheath gas flow rate at 35 units and auxiliary gas flow rate at 15 units. High resolution mass spectra were acquired as described previously.^40^

#### Metabolomic Analysis

For the metabolomic analysis the Metabolomics-SNET-V2 (release_23) workflow was used. Parameters were adapted from the GNPS documentation: MS2 spectra were filtered so that all MS/MS fragment ions within +/- 17 Da of the precursor *m/z* were removed. The MS/MS fragment ion tolerance and the precursor ion mass tolerance were set to 2.0 and 0.5 Da, respectively. Edges of the created molecular network were filtered to have a cosine score above 0.7 and at least 4 matched peaks between the connected nodes. The maximum size of molecular families in the network was set to 100. The MS2 spectra in the molecular network, filtered in the same manner as the input data, were searched against our internal library of 480 annotated metabolites. Reported matches between network and library spectra were required to have a score above 0.75 and at least 5 matching peaks. The molecular networks were visualized using Cytoscape.

#### Metabolite Quantification

Pseudouridine-containing intermediate were quantified by HPLC assuming an identical chromophore as PUM, against a purified pseudouridimycin internal standard. Relative amounts of the other metabolites were estimated as peak intensity ratio to those observed in WT strain.

#### Nucleotide sequence accession number and Paired Omics Data Platform project identifier

The genome sequence has been deposited in GenBank under the accession CP049782 as BioProject PRJNA609626. A subset of metabolomic data has been deposited in the Paired Omics Data Platform (Metabolomics project identifier c86fdc82-0d18-45d0-aa30-1f877c1cd3fc.2).

## ACKNOWLEDGMENTS

This work was supported by grants to Naicons from Regione Lombardia and Italian Ministry of Research (Nos. 30190679 and DM60066), from the European Union’s Horizon 2020 research and innovation program under grant agreement No.721484 (Train2Target), and from NIH (GM041376 and AI104660) to RHE.

## SUPPLEMENTARY FIGURES

**Supplementary Figure 1.**
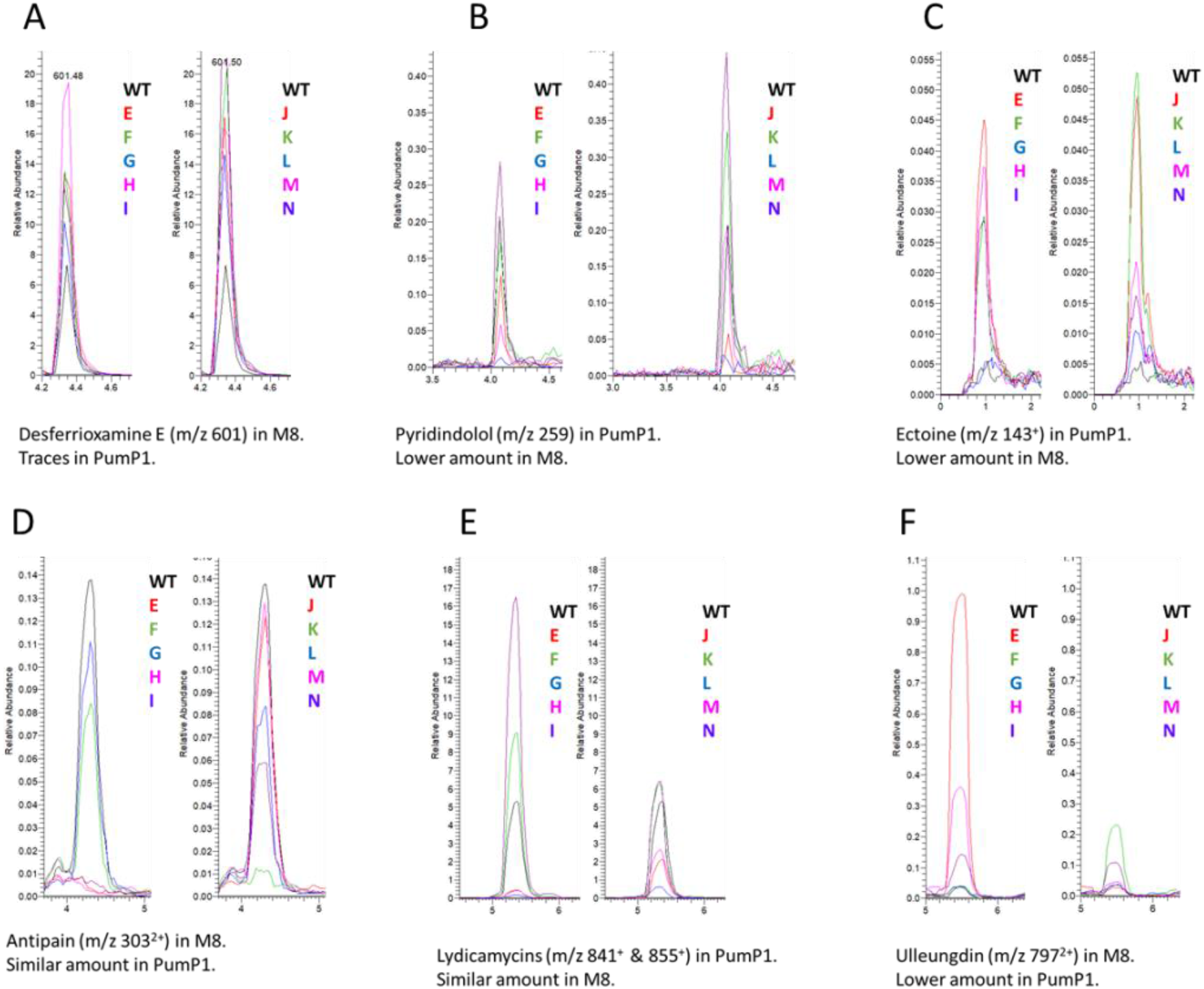
Comparison of extracted ion chromatograms related to desferrioxamine E(A), pyridindolol (B), ectoine (C), antipain (D), lydicamycin and ulleungdin (F) in the best medium from all the mutants and the wild type strain. Each comparison is split into two panels for clarity, with the WT added for correlation. *pum* mutants are designated by their letter suffix and color coded.

**Supplementary Figure 2.**
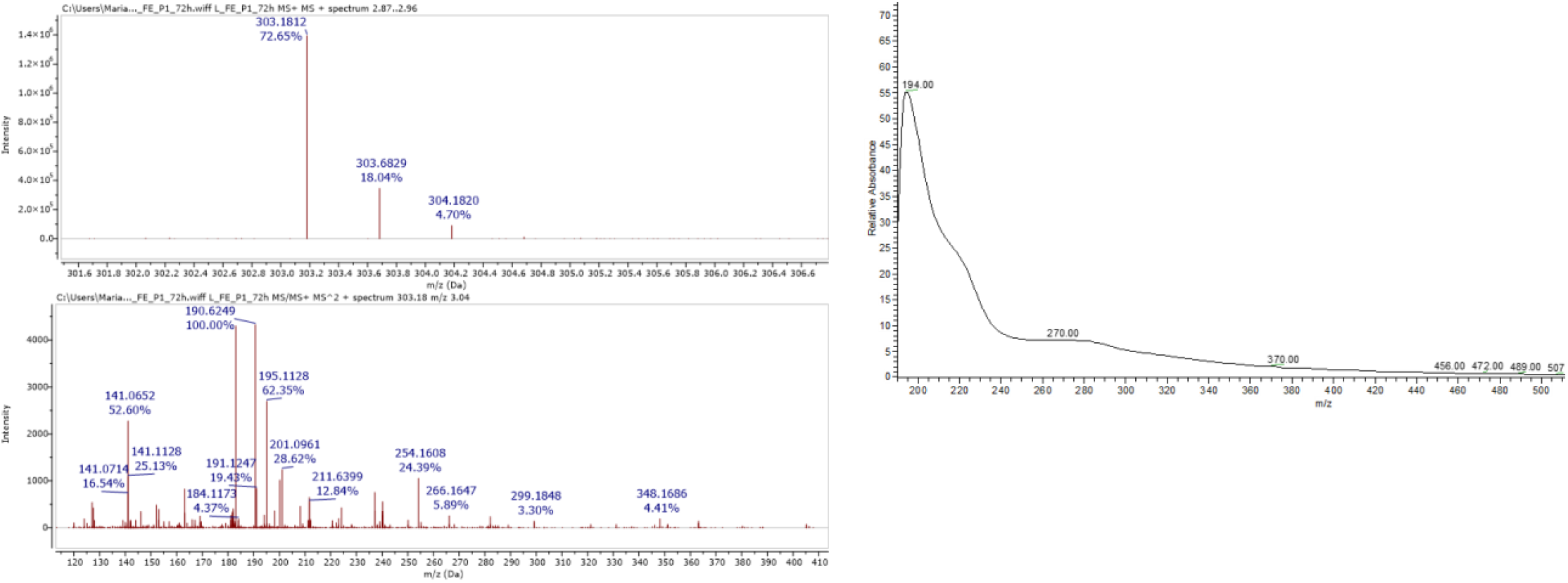
HR-MS and fragmentation spectrum of antipain and its UV-Vis spectrum.

**Supplementary Figure 3.**
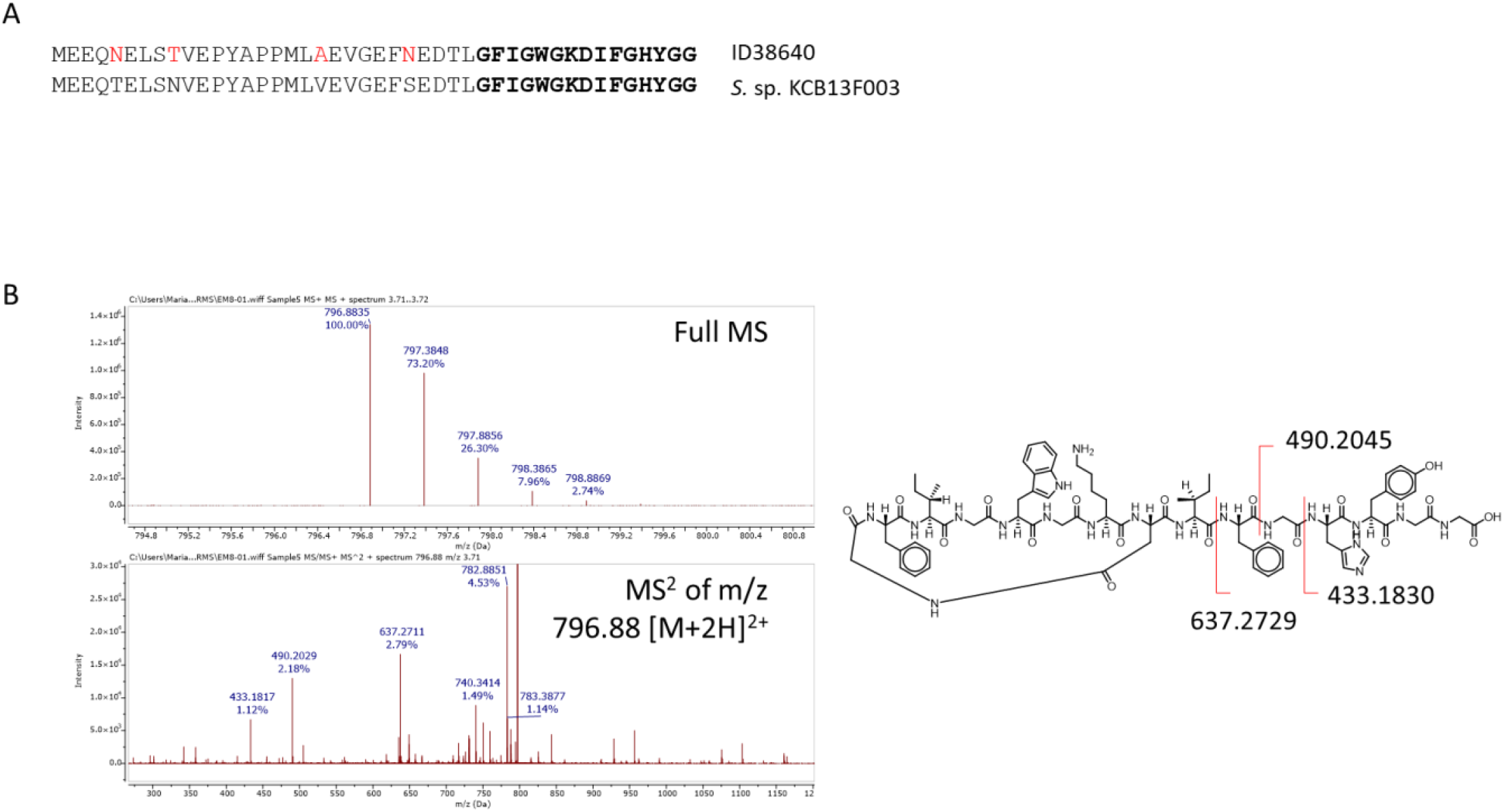
A) Precursor peptide of ulleungdin in ID38640 and in *S. sp*. KCB13F003. The different amino acids residues between the two leader sequences are highlighted in red type, while the core peptide is in bold type. B) HR-MS and fragmentation spectrum of ulleungdin (right side) and its annotated fragmentation pathway (left side).

**Supplementary Figure 4.**
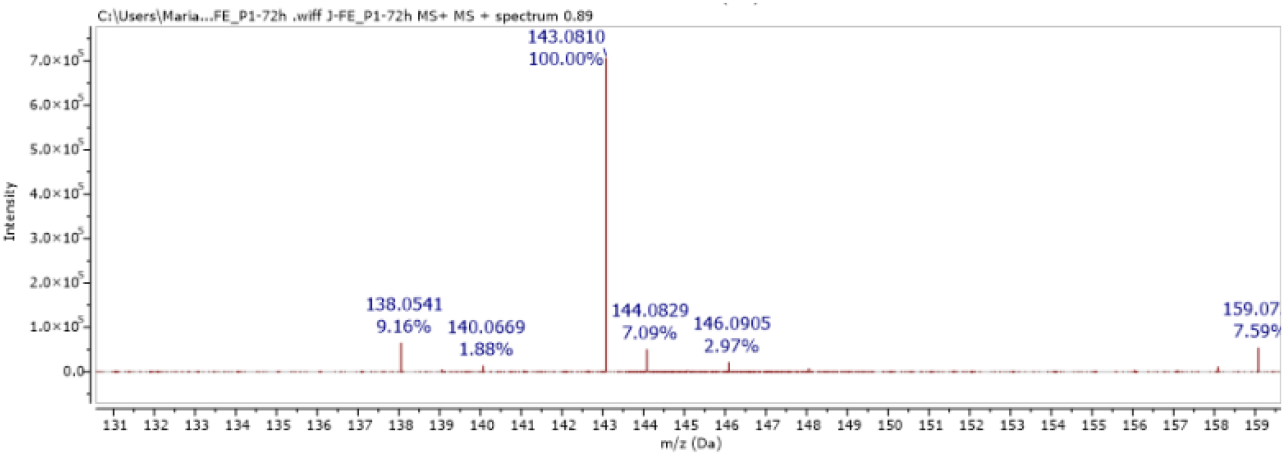
HR-MS of ectoine.

**Supplementary Figure 5.**
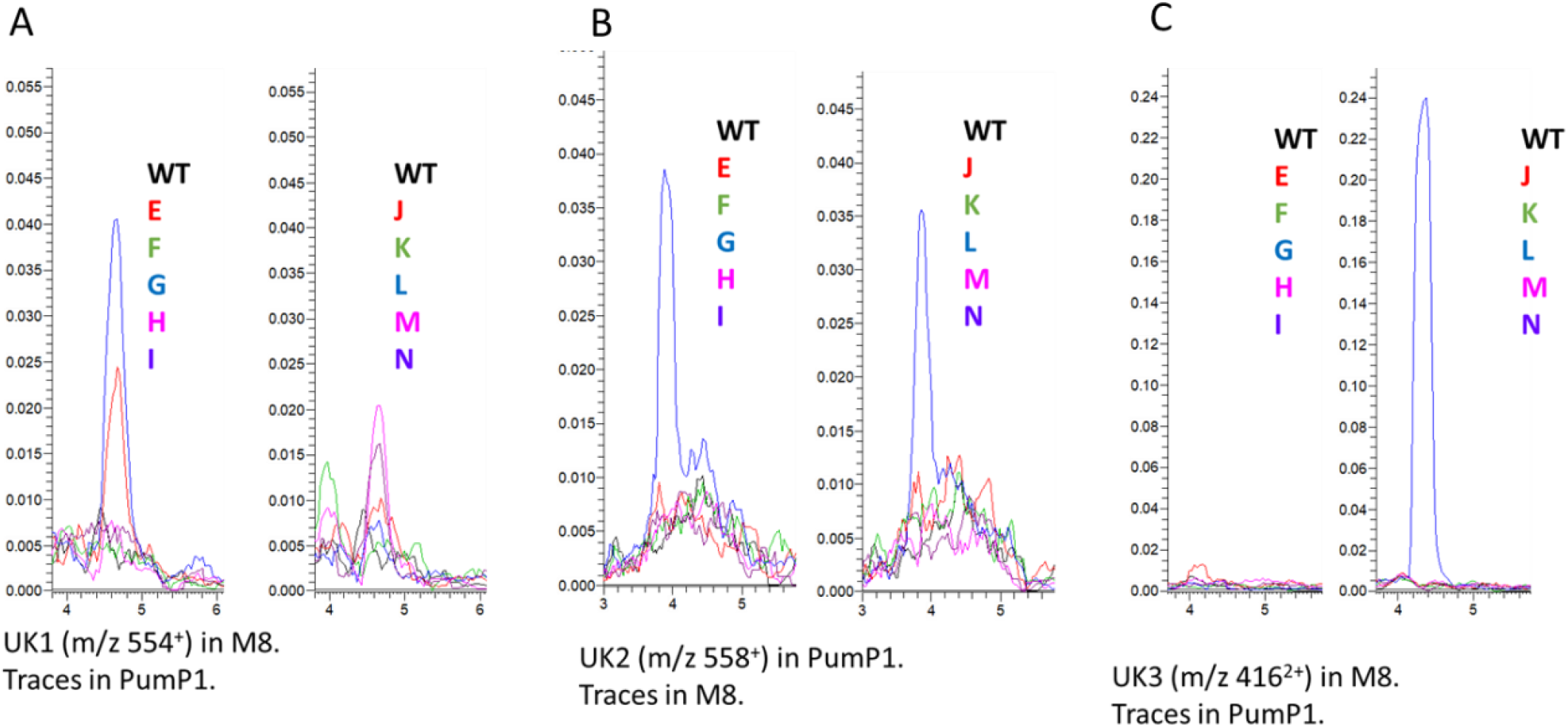
Comparison of extracted ion chromatograms related to the most representative species of the unannotated molecular families UK1 (A), UK2 (B) and UK3 (C) in the best medium from all the mutants and the wild type strain. Mutants labled as in Supplementary Fig. 1.

**Supplementary Figure 6.**
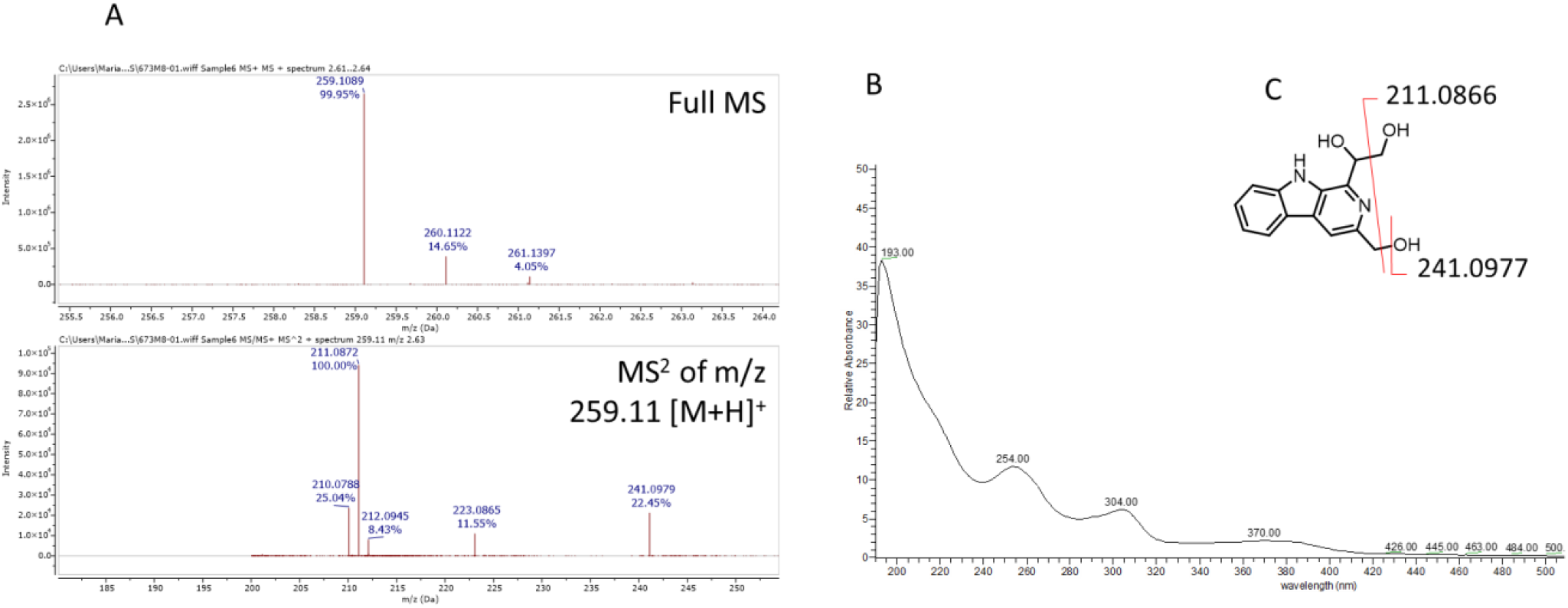
A) HR-MS and fragmentation spectrum of pyridindolol, its UV-Vis spectrum (B) and annotated fragmentation pathway (C).

**Supplementary Figure 7.**
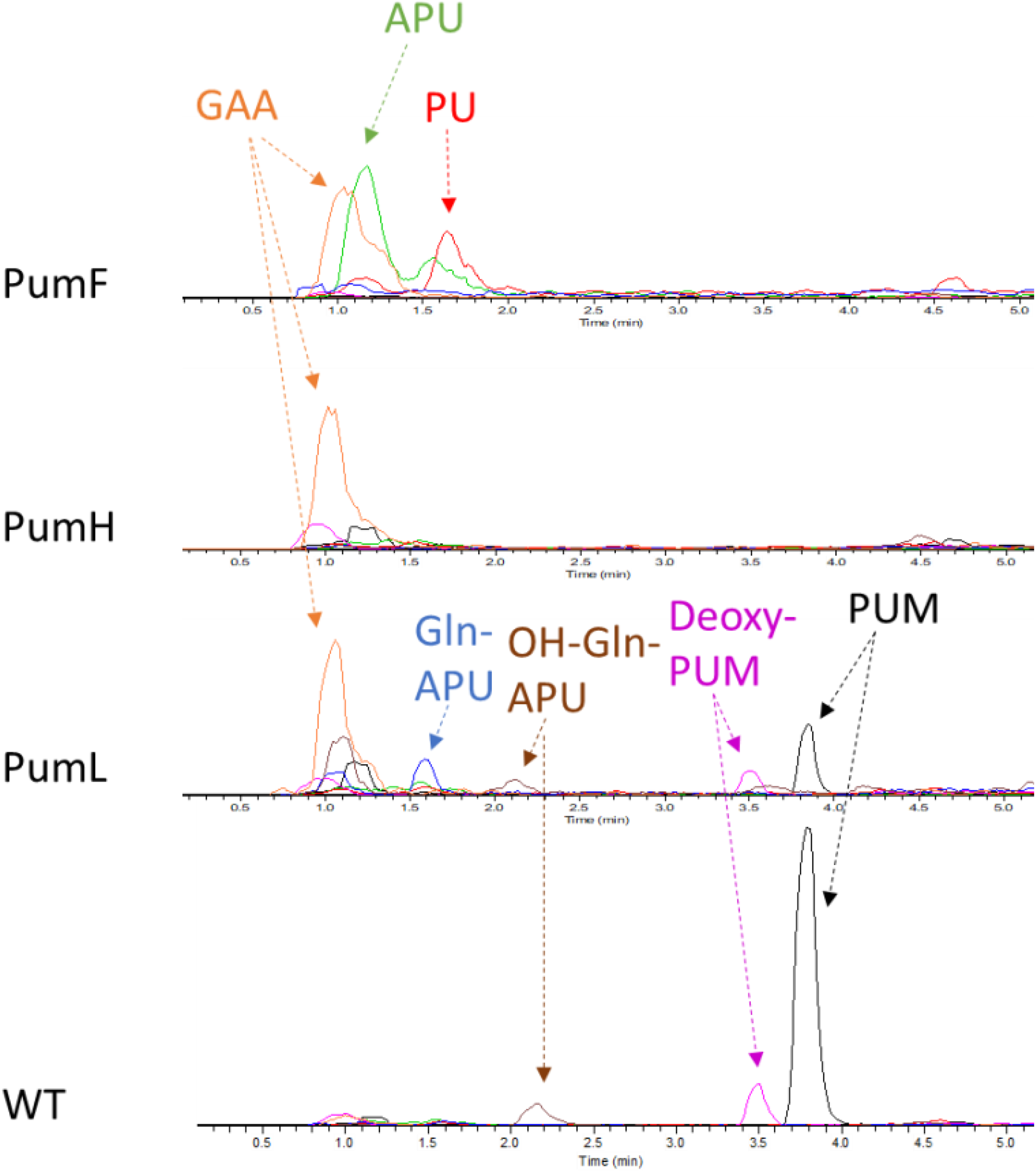
Extracted ion chromatograms of the *pumF, pumH* and *pumL* mutants. The analyses show pseudouridimycin (PUM, m/z 487 [M+H]+, black line), pseudouridine (PU, m/z 245 [M+H]+, red line), aminopseudouridine (APU, m/z 244 [M+H]+, green line), Gln-APU (m/z 372 [M+H]+, blue line), OH-Gln-APU (m/z 388 [M+H]+, brown line), deoxy-PUM (m/z 471 [M+H]+, pink line) and guanidinoacetic acid (GAA m/z 344 [M+2TFA-H]-, orange line).

**Supplementary Figure 8.**
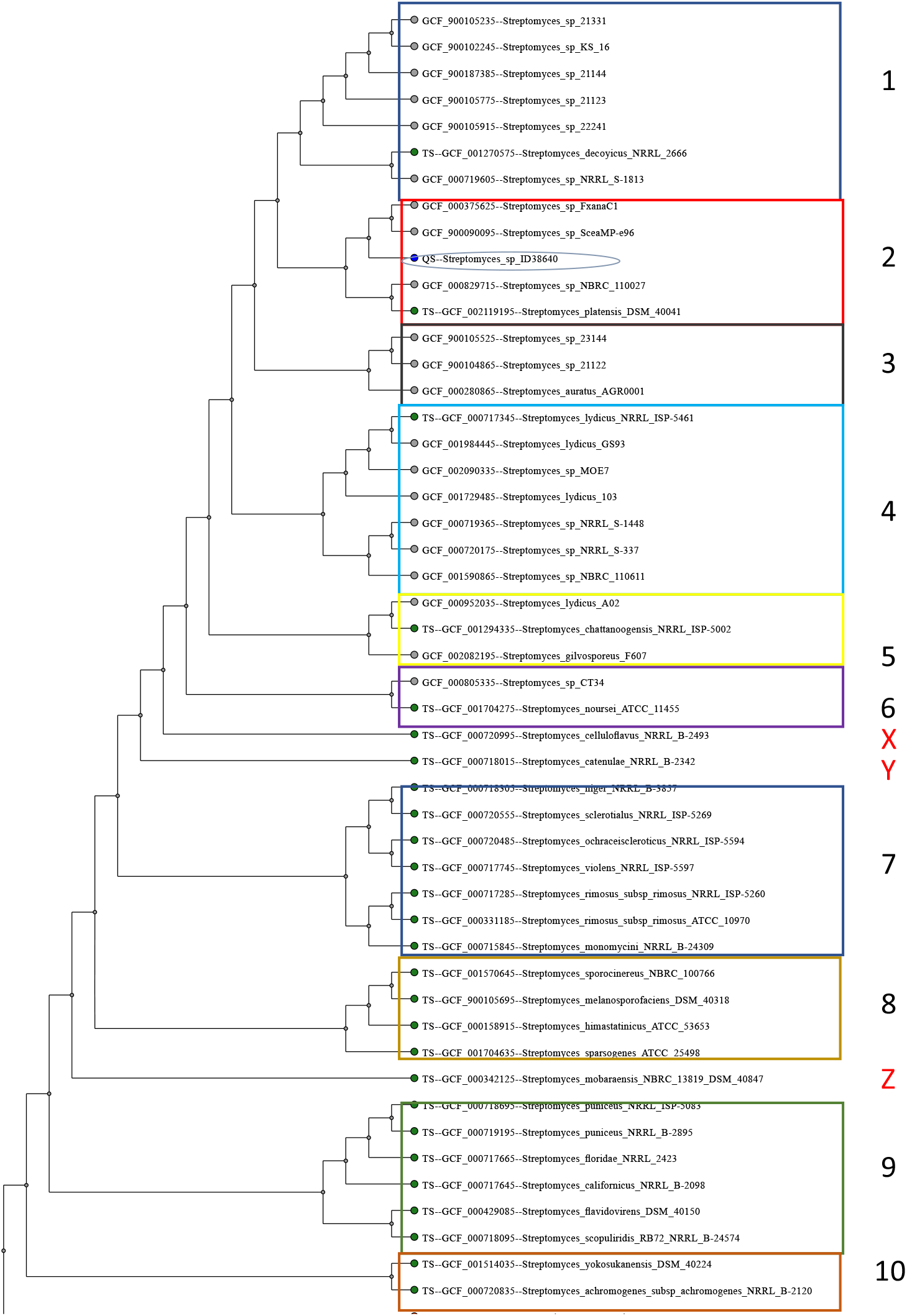
Maximum likelihood tree of 50 members of the genus *Streptomyces* based on 100 concatenated housekeeping genes identified by autoMLST. Color code is representing the nine major clades while X, Y and Z designated the three single strain branches. ID38640 is highlighted by a green circle oval.

**Supplementary Figure 9.**
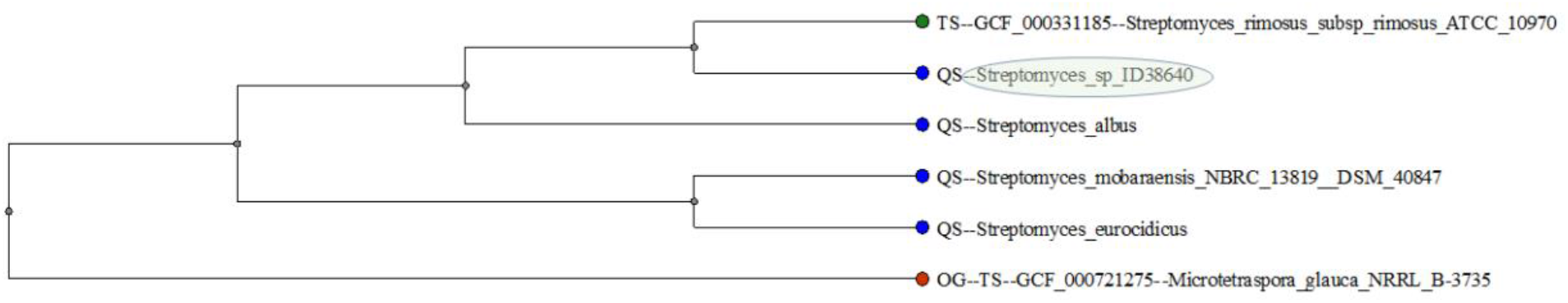
Maximum likelihood tree generated of ID38640, *S. eurocidicus, S. albus, S. mobaraensis*, and *S. rimosus* (*Microtetraspora glauca NRRL B-3735* used as the outgroup), based on 100 concatenated housekeeping genes identified by autoMLST. ID38640 is highlighted by a green circle.

**Supplementary Table 1.**
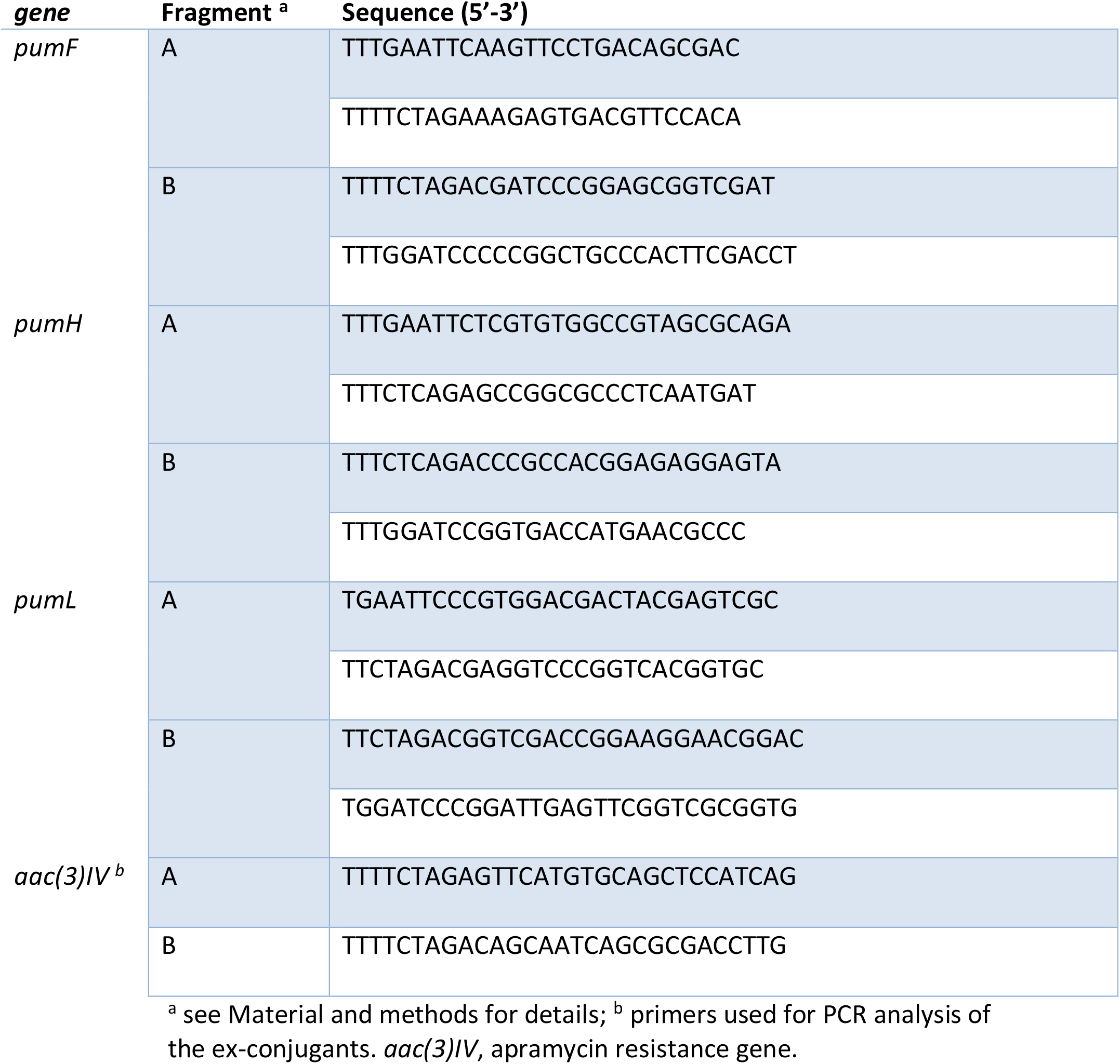
List of primers used for knock outs experiments.

## Notes

### Competing Interest Statement

The authors have declared no competing interest.

